# The grain amaranth pangenome reveals domestication-associated changes in diversity and function of structural variation

**DOI:** 10.64898/2026.01.08.698315

**Authors:** Ella Ludwig, Tom S Winkler, Markus G Stetter

## Abstract

**Background:** Grain amaranth is a nutritious pseudocereal from the Americas that was independently domesticated three times from a common wild ancestor. The three domesticated grain amaranths, their wild progenitor, and a close wild relative form a species complex. Pangenomes enable the assessment of genetic variation beyond single nucleotide polymorphisms.

**Results:** We have constructed a pangenome for the entire grain amaranth species complex, consisting of new, chromosome-scale genome assemblies for all five species, including the first reference genomes for *A. caudatus* and *A. quitensis*. Our high-quality assemblies reach near telomere-to-telomere contiguity. Comparative analyses within the grain amaranth pangenome revealed a high degree of collinearity and overall conserved chromosome structure across species. Genes are similarly conserved, with a ∼75% core gene set. We identify over 100,000 structural variants, distributed throughout the genomes. We quantify gene presence-absence and find that protein biosynthesis gene families were expanding during domestication, while gene loss reflects possible redundancies in other processes. We further map flowering time in a biparental population and find two QTL that together account for a 55-day difference in flowering time between homozygous genotypes. One QTL contains an ortholog of a known flowering-time regulator that may be disrupted by an insertion in the late flowering parent.

**Conclusions:** Our work establishes high-quality genomic resources for the promising protein crop grain amaranth and sheds light on how structural variants shape genomic diversity and repeated evolutionary change in crops. The structural variants and flowering time loci identified can help to understand amaranth adaptation and provide breeding targets for crop improvement.

## Background

Plant cultivation has been essential to human societies for millennia. During this long history of cultivation, selection has shaped plant traits to better meet human needs and preferences (Olsen and Wendel 2013; Purugganan 2019; Zohary 2004). This has led to a characteristic suite of morphological and physiological traits that distinguish cultivated crops from their wild ancestors, referred to as the domestication syndrome (Hammer 1984). The causal mutations responsible for these trait changes remain mostly unknown. However, evidence suggests that large scale genomic changes beyond single nucleotide mutations have often played an important role (Allaby 2014; Doebley et al. 2006). Uncovering the types of variation contributing to the genetic diversity of crops can help in understanding the process of plant adaptation and domestication.

Early studies of genetic variation and domestication relied on small genetic markers such as simple sequence repeats (SSRs, also known as microsatellites) and single nucleotide polymorphisms (SNPs). The development of high-throughput sequencing technologies enabled genome-wide identification of SNPs, which became the predominant marker type for population genetics and genotype-phenotype associations (Aoun et al. 2016; Moragues et al. 2010; Xiao et al. 2017). Structural variation, on the other hand, encompasses larger genetic polymorphisms, such as insertions, deletions, duplications, translocations, and inversions longer than 50 base pairs (Ho et al. 2020). Notably, at least one-third of known domestication loci in plants are linked to structural variants (SVs), highlighting their evolutionary importance (Gaut et al. 2018). Beyond their role in domestication and impact on phenotypes, SVs contribute a substantial amount of genetic variation in a crop (Zhou et al. 2019). In fact, increasing evidence suggests that SVs may explain more phenotypic variation than SNPs (Chaisson et al. 2019; Gaut et al. 2018; Lye and Purugganan 2019; Yang et al. 2019; Zhou et al. 2019). Comparative studies have identified thousands of SVs, both across species (Stein et al. 2018; Zhao et al. 2018), and even between individuals of the same species (Brunner et al. 2005; Gordon et al. 2017; Maretty et al. 2017; Sun et al. 2018). For example, a comparison between just two maize inbred lines revealed that more than 20% of genes contained large SVs or other large-effect mutations (Sun et al. 2018), highlighting SVs as an essential component of genetic diversity (Chaisson et al. 2019; Gaut et al. 2018; Zhou et al. 2019). Given their potential to shape key traits and genetic variation, a deeper understanding of SVs and their role in plant domestication and crop diversity is essential.

Despite their importance, SVs have historically been understudied, largely because earlier sequencing technologies made it difficult to assemble genomes with structural variation accurately resolved (Lemay and Malle 2022). Only with the advent of third generation sequencing technologies, in particular the development of highly accurate long reads, has genome assembly quality improved enough to reliably characterize SVs (Garg et al. 2024; Jiao and Schneeberger 2017). Still, progress has been constrained not only by incomplete assemblies but also by the continued reliance on single reference genomes, which — even when high quality — cannot capture the full spectrum of structural diversity within a species (Ahmad et al. 2025; Garrison *et al*. 2018; Secomandi et al. 2025; Yang et al. 2019). Pangenomes have emerged as a more comprehensive framework for capturing structural genomic diversity (Bayer et al. 2020; Du et al. 2025; He et al. 2025). They integrate sequences from multiple representative individuals, and thereby capture variation present within a species and beyond (Matthews et al. 2024; Schreiber et al. 2024). Many existing plant pangenome studies focus on a single species and are primarily limited to cultivated accessions (Khan et al. 2020). However, crop wild relatives harbor a vast amount of unexplored genetic diversity that cultivated accessions have lost during domestication and subsequent breeding (Khan et al. 2020; Schreiber et al. 2024). This often includes genes underlying disease resistance or tolerance to abiotic stresses — traits that are valuable for crop improvement and can serve as direct targets for breeders (Hu et al. 2025; Schreiber et al. 2024). Including crop wild relatives in pangenomes is therefore essential for fully understanding the genomic basis of domestication, particularly the role of SVs in shaping this process.

Pangenomes are starting to become the standard for main crops, but for non-model crops, a good representation of genomic diversity is often still missing. However, these crops and their wild relatives harbor particularly high diversity and have high potential for future agricultural systems (Khan et al. 2020; Schreiber et al. 2024; Lara 2025). The grain amaranths (*Amaranthus* spp.) are highly diverse and include three domesticated species (Stetter et al. 2020). Amaranth is a highly nutritious pseudocereal native to the Americas, where it served as a major staple crop prior to European colonization (Sauer 1967). Its seeds are gluten-free and rich in protein, fiber, and essential micronutrients such as iron and calcium (Angel Huerta-Ocampo and Paulina Barba de la Rosa 2011; Rastogi and Shukla 2013). Beyond their nutritional value, grain amaranths are resilient under abiotic stresses, including drought, heat, and salinity (Barba de la Rosa et al. 2009; Joshi et al. 2018; Pulvento *et al*. 2022), characteristics that make them increasingly relevant in the face of climate change. The grain amaranth complex includes five species: the three domesticated species (*Amaranthus cruentus* L., *Amaranthus hypochondriacus* L., and *Amaranthus caudatus* L.) and two wild relatives (*Amaranthus hybridus* L. and *Amaranthus quitensis* Kunth.; Figure 1A). All three domesticates were most likely domesticated from *A. hybridus*, while *A. quitensis* may represent either a wild intermediate between *A. hybridus* and *A. caudatus* (Gonçalves-Dias et al. 2023) or feral type of *A. caudatus* (Figure 1A; Stetter and Schmid 2017). The five diploid species, with an estimated genome size of ∼500 Mb, are closely related and shared extensive gene flow after domestication (Gonçalves-Dias et al. 2023). While a high-quality reference genome for *A. hypochondriacus* (Graf et al. 2025) and short-read whole-genome sequencing of hundreds of individuals (Stetter et al. 2020; Singh and Stetter 2025) have shown the high diversity of the crops and their wild relatives, the diversity of structural variation remains unknown.

**Figure 1.**
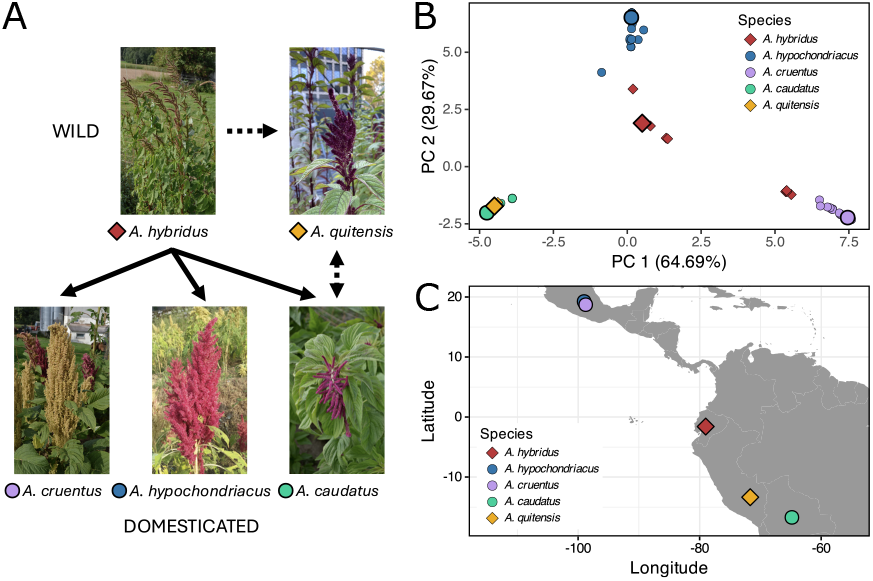
Relationships among the five grain amaranth species and the genetic and geographic distribution of the sequenced accessions. (A) Schematic representation of grain amaranth domestication, with photographs of representative plants. Arrows indicate the direction of gene flow during domestication, with dashed arrows representing relationships that are not yet resolved. Colored diamonds next to species names indicate wild species, while colored circles indicate domesticated species. (B) PCA of 88 individuals showing the genetic distribution across species in the grain amaranth complex, with individuals sequenced in this study shown as larger points. Points are colored by species and shaped to distinguish wild from domesticated accessions, as in A. (C) Map showing the geographic origins of the five newly sequenced accessions, colored and shaped as in A and B.

In this study, we generate a pangenome for the grain amaranth species complex — including the three domesticated grain amaranths, their common wild ancestor, and a closely related wild species — to investigate the genomic changes underlying repeated domestication. We assembled and annotated all five species using a unified workflow and achieved high-quality nuclear, chloroplast, and mitochondrial genomes. These genomic resources enable direct comparison of genome structure, synteny, and variation across the species complex. We distinguished conserved and lineage-specific genes and examined gene gain and loss in domestication. We find gain and loss in genes that perform central cellular functions, pointing to functional redundancies, and underscoring the polygenic nature of domestication and the diversity within this crop. We find two major quantitative trait loci (QTL) controlling flowering time in amaranth, using linkage mapping in over 400 wholegenome-sequenced individuals. An SV within one of the QTL is a potential causal mutation disrupting gene function and delaying flowering by 11.5 days.

## Results

### Genome assemblies

Five grain amaranth accessions, selected as representative examples of their respective species from their native range (Figure 1), were sequenced with PacBio HiFi long-read sequencing. Across all five species, sequencing reads show both high per-base accuracy (mean QV 24.4-30.0) and long average read lengths (∼12 kb or longer; Table S1) with a sequencing depth ranged from 33x to 68x (Table S1).

We assembled reference genomes for the whole grain amaranth species complex, including the three domesticates and two wild ancestors using Hifiasm and scaffolded the primary contigs against the published *A. hypochondriacus* v3 reference genome without breaking assembled contigs. The assemblies were chromosome-scale, representing the 16 *A. hypochondriacus* chromosomes. The genome assembly lengths varied from 415.4 Mb in *A. hybridus* to 435.2 Mb in *A. hypochondriacus* (Table 1). These sizes are comparable to the *A. hypochondriacus* v3 reference genome (434.9 Mb), with the new *A. hypochondriacus* assembly differing by less than 1 Mb in length from the reference, suggesting little within-species variation in genome size (Stetter and Schmid 2017).

**Table 1.** Genome assembly and annotation statistics for five new grain amaranth assemblies compared to the published *A. hypochondriacus* v3 reference (Graf et al. 2025). Estimated genome sizes are from Stetter and Schmid (2017).

|  | *A. hypochondriacus*<br>v3 reference | *A. hybridus* | *A. hypochondriacus* | *A. cruentus* | *A. caudatus* | *A. quitensis* |
| --- | --- | --- | --- | --- | --- | --- |
| Est. genome size (Mb) | 506.4 | 503.8 | 506.4 | 510.9 | 502.0 | 501.1 |
| Assembly length (Mb) | 434.9 | 415.4 | 435.2 | 432.5 | 416.4 | 431.1 |
| Number of scaffolds | 232 | 91 | 202 | 140 | 120 | 253 |
| Portion of assembly<br>anchored to chromosomes | 96.5% | 98.3% | 96.5% | 96.5% | 97.7% | 95.5% |
| Contig N50 (Mb) | 9.6 | 3.2 | 4.6 | 3.3 | 2.7 | 1.8 |
| Scaffold N50 (Mb) | 26.0 | 24.4 | 25.5 | 26.8 | 24.6 | 25.4 |
| LTR Assembly Index (LAI) | 16.32 | 17.32 | 16.79 | 16.60 | 18.33 | 16.99 |
| Quality Value (QV) scores | 51.9 | 61.0 | 64.1 | 61.0 | 59.5 | 60.5 |
| K-mer completeness | 97.0% | 98.4% | 98.7% | 98.5% | 98.5% | 99.5% |
| BUSCO score (proteome) | 98.8% | 97.1% | 97.3% | 96.9% | 97.3% | 97.3% |
| Number of genes | 25,167 | 25,849 | 27,632 | 25,562 | 24,944 | 25,567 |
| Total TE content | 50.4% | 47.9% | 49.3% | 51.6% | 51.1% | 51.2% |

The total proportion of TEs ranged from 47.9% in *A. hybridus* to 51.6% in *A. cruentus* (Table 1). Predicted gene counts from Helixer varied between 24,944 in *A. caudatus* and 27,632 in *A. hypochondriacus*, compared to 25,167 genes in the reference (Graf et al. 2025), but similar to the closely related sugar beet (27,421; Dohm et al. 2014) and spinach (28,247; She et al. 2025).

Overall, genome quality was as high or higher in the new assemblies compared to the high-quality *A. hypochondriacus* v3 reference genome of PI 558499 (Graf et al. 2025). The number of scaffolds was lower in all but one assembly, and the scaffold N50 was over 24 Mb in all assemblies, indicating very contiguous assemblies. Importantly, more than 95.5% of the assembly length in each species was able to be anchored to chromosomes, and this proportion was higher than the reference genome in all but one assembly. The LTR Assembly Index (LAI) also showed that these assemblies are all highly contiguous, with LAI scores between 16.60 and 18.33 and K-mer-based analysis showed quality value (QV) scores ranging from 59.5 to 64.1 (Table S1), corresponding to base-level accuracies of ∼99.9999%, or roughly only one error per 900,000 bases (Rhie et al. 2020). K-mer-based completeness values ranged from 98.4% in *A. hybridus* to 99.5% in *A. quitensis*, demonstrating that nearly all k-mers from the read sets were represented in the assemblies (Table 1). BUSCO analysis of protein-coding genes returned scores 96.9% or above for all assemblies, indicating their high completeness based on gene content. Overall, these results establish that the five new genomes are highly contiguous, complete, and correct, laying the foundation for the grain amaranth species complex pangenome.

### Grain amaranth pangenome reveals a highly conserved genomic landscape

The chromosomal distribution of genes and TEs provides insight into the structural organization of genomes, particularly with respect to centromeres and telomeres. Across all five genomes, chromosomes exhibited an uneven distribution of genes and TEs (Figure S1), with the expected inverse relationship between gene and TE density observed in plants (Naish et al. 2021; Ren et al. 2023; Sidhu and Gill 2005; Sun et al. 2018; Yang et al. 2024) and previously reported in *A. hypochondriacus* (Graf et al. 2025). The centromere annotation revealed that chromosome 1-6, 12, and 15 are metacentric, while 7-11, 13, and 16 appear telocentric. The distribution of TEs and genes follows this pattern, confirming the centromere annotation (Figure S1). Two exceptions were observed: chromosome 16 in *A. caudatus* and chromosome 6 in *A. quitensis*, where the centromeres were predicted in regions that were gene-rich and TE-poor. Centromere repositioning has been documented in soybean (*Glycine max*) via structural variation and bursts of TE activity (Liu et al. 2023), raising the possibility that chromosomes 16 in *A. caudatus* and 6 in *A. quitensis* represent recent centromere shift, which could account for the atypical gene and TE density patterns at the centromere. The patterns suggest active genome evolution in the relatively young species complex.

We also annotated telomeres, which mark chromosome ends and indicate genome completeness. Telomeres were identified by searching for the canonical telomeric repeat “AAACCCT.” In all five assemblies, every chromosome contained at least one terminal telomere, and on average 63.5% of chromosomes per assembly contained telomeres at both ends, highlighting the near telomere-to-telomere (T2T) resolution of our assemblies (Figure S2; Table S2). Three chromosomes — chromosomes 1 and 7 in *A. cruentus* and 7 in *A. caudatus* — contained three telomeric repeat arrays, suggesting potential misassemblies, as only two telomeres are expected per chromosome. Overall, the centromeres and telomeres provide further evidence for the high completeness of our assemblies, with multiple chromosomes achieving T2T resolution.

In addition to nuclear assemblies, we generated complete chloroplast and mitochondrial assemblies for each accession, enabling cross-species comparisons of organellar genome size, gene content, and structural organization. All chloroplast genomes assembled into a single circular contig with the canonical quadripartite structure, consisting of a large single-copy (LSC) region of ∼85.6-85.8 kb, a small single-copy (SSC) region of ∼17.95-17.97 kb, and two inverted repeats (IRs) of ∼23.53 kb (Figure S3; Table S3). Total chloroplast genome sizes were highly similar, ranging from 150.6 to 150.8 kb, with only 164 bp difference between the smallest (*A. caudatus*) and largest (*A. hybridus*) assemblies. Each chloroplast genome contained 126-127 unique genes, including 79 protein-coding genes, 39-40 tRNAs, and 8 rRNAs. In contrast to the chloroplast, mitochondrial genome sizes showed greater variation, ranging from 339.8 kb in *A. cruentus* to 419.2 kb in *A. caudatus* (Figure S4; Table S3). Despite this size variation, GC content was consistent across species (42.6-42.8%), and gene content similarly conserved. Each mitochondrial genome contained 61-62 unique genes, including 33-34 protein-coding genes, 25 tRNAs, and 3 mRNAs. *A. hybridus* and *A. caudatus* each lacked one protein-coding gene (*ccmB*) present in the other three species. Together, the nuclear, chloroplast, and mitochondrial assemblies provide near-complete genomes for each species, enabling robust analyses of domestication history across the species complex.

### Structural variation highlights genomic diversity across grain amaranth species

To assess the large-scale structural variation of the grain amaranth species complex, we evaluated the nuclear genome synteny at both a whole-genome and chromosome-by-chromosome scale across the five species. The analysis revealed that largescale chromosomal structure was highly conserved across all species, with most orthologous regions corresponding to homologous chromosomes, showing a clear one-to-one chromosome correspondence (Figure 2). This conserved structure was consistent across both wild and domesticated species. Many chromosomes, such as 1, 11, and 16, were almost completely conserved across species, showing few breaks in large-scale syntenic blocks. By contrast, chromosomes 2, 3, 4, 9, and 13 showed larger gaps in synteny or more extensive rearrangements between some species, suggesting increased large-scale structural divergence among these homologous chromosomes (Figure 2). Large-scale intrachromosomal rearrangements were also evident on chromosome 15 in *A. caudatus* and *A. quitensis*. In *A. caudatus*, there are several larger translocations and inversions relative to *A. cruentus*, whereas this entire region is one large inversion in *A. quitensis*.

**Figure 2.**
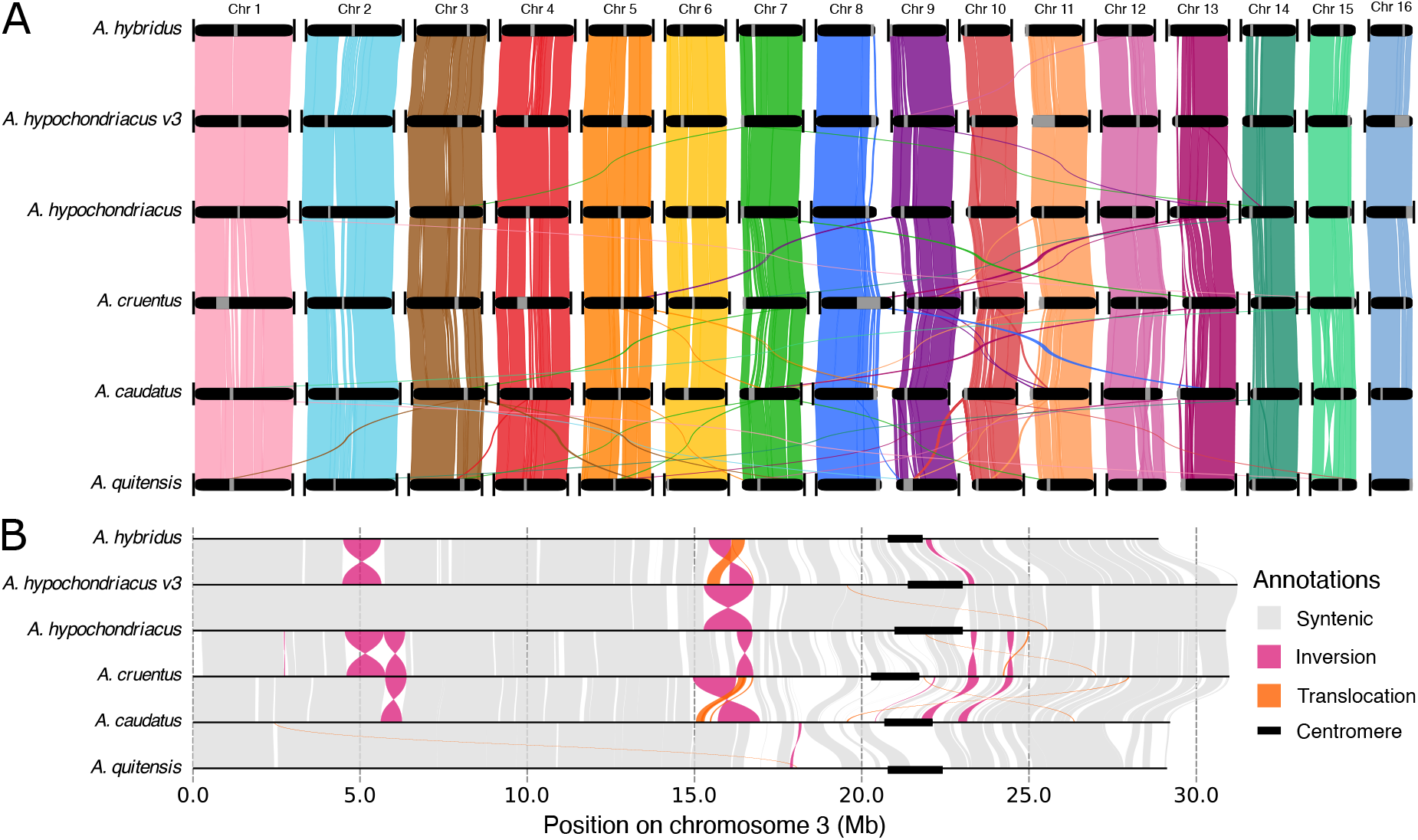
Synteny among six grain amaranth genome assemblies. (A) Whole-genome synteny based on DNA sequence. Chromosomes are ordered numerically from left to right; gray bands denote centromeres, and vertical lines indicate telomeres at the ends of chromosomes. Ribbons representing syntenic regions are colored according to their corresponding regions in the assembly above. (B) Intrachromosomal synteny of chromosome 3. Syntenic blocks shown in gray, inversions in pink, translocations in orange, and centromeres are annotated with a black box.

Within chromosomes, additional SVs become apparent (Figure 2B; Figure S5). For instance, on chromosome 3, SVs were primarily concentrated in three regions, at ∼5 Mb, at ∼16 Mb, and at ∼23 Mb. An inversion centered at ∼5 Mb in *A. hypochondriacus* relative to *A. hybridus* is not shared by the other species, and a second inversion at ∼6 Mb is unique to *A. cruentus*. Together, these features illustrate both the broad conservation of chromosome structure across species and the lineage-specific rearrangements that contribute to structural diversity within the complex.

While visual synteny illustrates large scale SVs, it is difficult to compare these between species. To understand the genome evolution and diversity related to the domestication of amaranth, we called SVs against an outgroup species. We called SVs using SVIM-asm (Heller and Vingron 2021) relative to the *A. retroflexus* reference genome (Raiyemo et al. 2025) which is equally diverged from all six (including the *A. hypochondriacus* reference genome) species. This identified 105,702 unique SVs, of which 96.7% were insertions and deletions (Figure 3A & B). The distributions of insertions, deletions, and duplications were skewed towards smaller variants with lengths less than 10 kb, while inversions were almost all longer than 10 kb (Figure 3A). All six genome assemblies contained SVs of all types, with duplications being the least common type across species (Figure 3B). The total number of SVs per assembly ranged from 39,418 in *A. hybridus* to 41,207 in *A. quitensis*.

**Figure 3.**
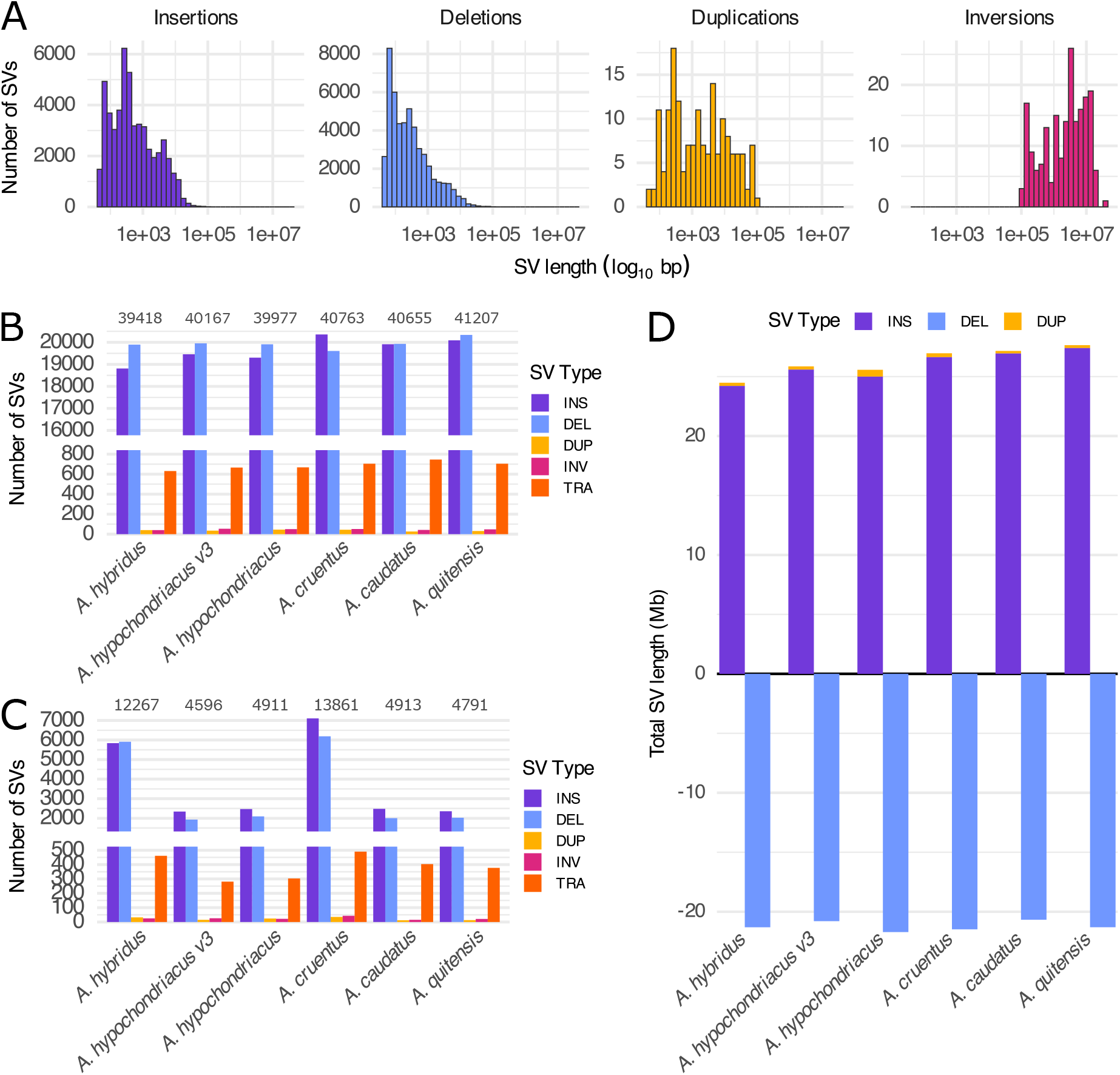
Structural variants (SVs) identified in six grain amaranth genomes relative to the *A. retroflexus* reference genome (Raiyemo et al. 2025). (A) Histograms of SV lengths in bp (x-axis is in log_10_ scale). Plots are faceted and colored by SV type: insertions (INS) in purple, deletions (DEL) in blue, duplications (DUP) in yellow, and inversions (INV) in pink. (B) Number of total SVs and (C) number of unique SVs detected in each of the six genomes. Bars are colored by SV type as in (A), with translocations (TRA) additionally colored in orange, and per-genome totals above bars. (D) Total length of assemblies contained in SVs that add or remove sequence from the genome. Bars show the sum of SV lengths (in Mb), with insertions (INS) and duplications (DUP) stacked above the x-axis and deletions (DEL) below the x-axis.

Comparing the number of unique SVs per species revealed differences between genomes. *A. hybridus* and *A. cruentus* had roughly three times as many unique SVs as the other four assemblies. The elevated number of unique SVs in *A. hybridus* is consistent with its higher genetic diversity as the wild ancestor of the domesticated species and increased outcrossing rate compared to the other species (Singh and Stetter 2025; Jain et al. 1982; Hauptli and Jain 1985; Nyambo et al. 2024). The two *A. hypochondriacus* genomes shared a large fraction of SVs, which led to fewer unique variants. Similarly, *A. caudatus* and *A. quitensis* shared many SVs, in accordance with their close genetic relationship, likely due to a shared domestication history (Stetter and Schmid 2017). The cumulative SV length of sequence-adding variants (insertions and duplications) together accounted for between 24.9 Mb in *A. hybridus* to 28.1 Mb in *A. quitensis*, whereas deletions summed to between 21.0 Mb in *A. caudatus* to 22.1 Mb in *A. hypochondriacus* (Figure 3D). Although deletions were more numerous than insertions in all genomes except *A. cruentus*, they consistently affected a smaller cumulative sequence length than insertions. Notably, although *A. hypochondriacus* did not have more duplications, it had a larger total duplication length than the other assemblies. These differences in cumulative SV length point to lineage-specific structural changes that could reflect the evolutionary and domestication histories among the grain amaranth species complex.

### Core gene content is highly conserved across the grain amaranths, with domestication likely reshaping existing pathways

An additional level of pangenomic diversity is gene content. To assess patterns of gene conservation and divergence among the five grain amaranth species, we identified orthologous genes, grouped into orthogroups, using OrthoFinder (Emms and Kelly 2019; Emms et al. 2025), and examined their distribution across the genomes. A total of 25,205 orthogroups were identified Table S4. Of these, 18,888 contained at least one gene of each of the six genomes (Figure 4A), constituting the core gene set and representing 74.9% of all orthogroups. The remaining 6,317 orthogroups (25.1%) were present in only a subset of the genomes and comprise the variable gene set (Figure 4A & B). Assembly-specific orthogroups were rare, with only 363 orthogroups (1.4%) unique to a single assembly. The number of assembly-specific orthogroups ranged from 5 in *A. caudatus* to 156 in the *A. hypochondriacus* v3 reference, while orthogroups shared by two to four assemblies ranged from 12 to 376 (Figure 4B).

**Figure 4.**
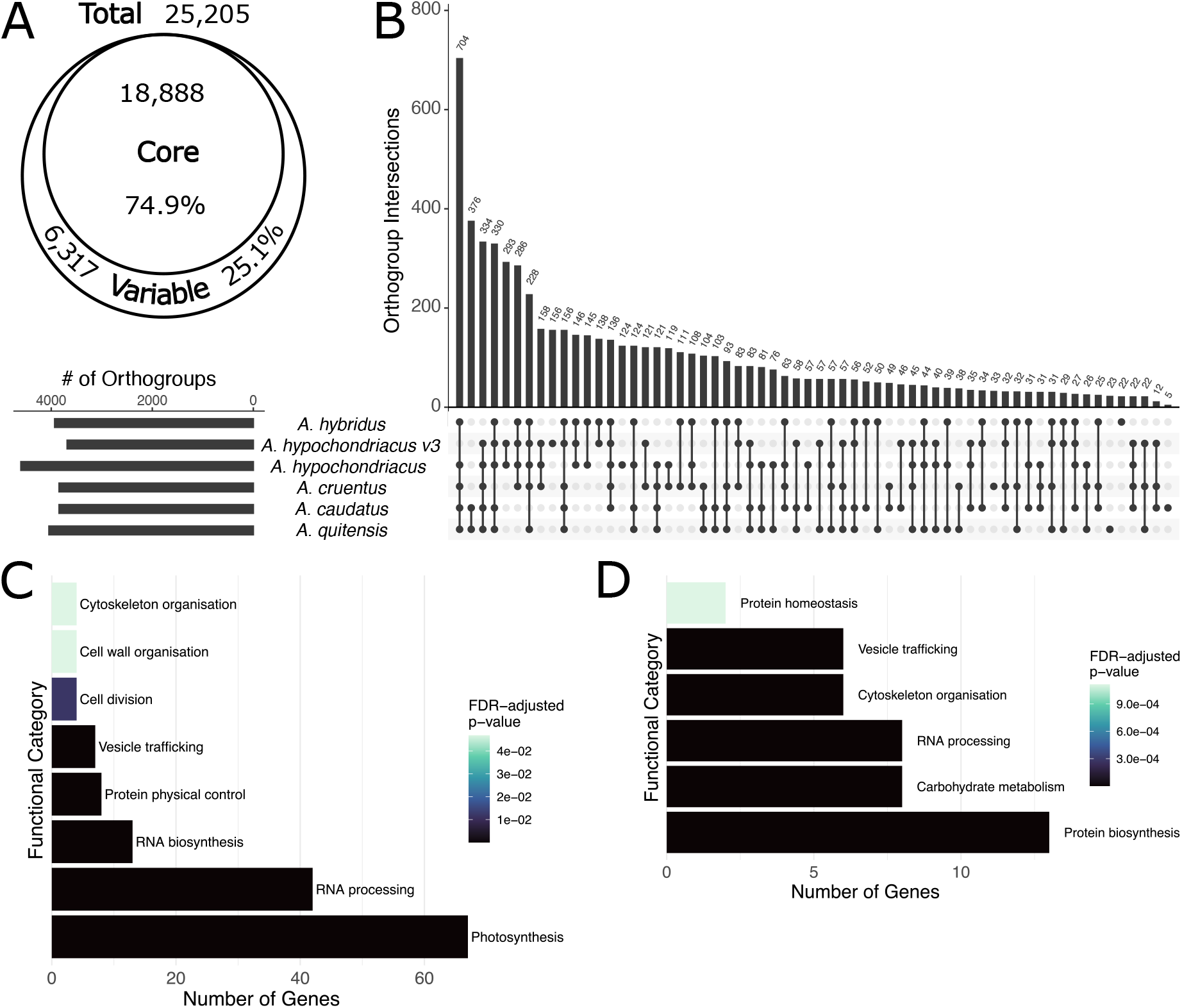
Orthologous gene relationships among six grain amaranth genome assemblies. (A) Diagram showing the number of core orthogroups (present in all six genomes) and variable orthogroups (present in only a subset of genomes). (B) UpSet plot of variable orthogroups identified by OrthoFinder, showing intersections between genomes and the number of genome-specific orthogroups. The horizontal barplot on the left indicates the number of orthogroups per genome, and the vertical bars show the number of orthogroups shared by the genome combinations indicated by the dots below. (C-D) Functional enrichment analysis of orthogroups containing more genes in the (C) wild ancestor *A. hybridus* compared to all domesticated species and (D) in the three domesticated species compared to *A. hybridus*, with MapMan categories colored by significance in the Mercator4 enrichment analysis (FDR-adjusted p-value).

We sought to understand the influence of domestication on the pangenome of amaranth. In other crops, domestication was accompanied by gene gain and loss (Domazet-Lošo et al. 2024; Clark 2023; Bayer et al. 2022). To investigate this, we examined the core orthogroups for changes in gene number in the domesticated species (*A. hypochondriacus, A. cruentus*, and *A. caudatus*) relative to their wild ancestor (*A. hybridus*). We identified 185 orthogroups that lost genes in all domesticated species and five orthogroups that gained genes. Functional enrichment analysis revealed that orthogroups losing genes in the domesticated species relative to *A. hybridus* were significantly enriched (FDR-adjusted p-value <0.05) across eight categories (Figure 4C), with the largest enrichment in photosynthesis-related genes. Orthogroups that gained genes were significantly enriched in six functional categories (Figure 4D), with protein biosynthesis most enriched.

A family of genes commonly lost or gained during domestication are nucleotide-binding leucine-rich repeat (NLR) genes (Barragan and Weigel 2021). This highly diverse gene family plays a central role in plant defense against pathogens (van Wersch and Li 2019) and copy number is often highly variable (Barragan and Weigel 2021). In grain amaranth, the total number of NLR genes was relatively conserved, with little variation of gene counts between wild and domesticated taxa. We found fewer NLR genes were predicted in *A. caudatus* and *A. quitensis* (164 and 175, respectively) — the two South American species — compared to the over 200 genes predicted in the two North American species and *A. hybridus* (Figure S6, Table S5).

In summary, gene gain and loss during grain amaranth domestication involved similar core functional categories, with gene loss being more prevalent, suggesting redundancy in gene function. Yet, the enrichment of protein biosynthesis and carbohydrate metabolism genes being gained might be related to the nutritional value of the crop.

### Mapping flowering time in A. hypochondriacus

Pangenome analysis has revealed the high diversity of SVs in the amaranth species complex. Often, genetic diversity contributes to phenotypic diversity. Flowering time is an essential live history trait in plants, that is also of utmost importance for crop yield. We performed QTL mapping for flowering time using recombinant inbred lines (RILs) in a bi-parental *A. hypochondriacus* mapping population (Winkler et al. 2026). We recorded the flowering time of the parental lines and 449 RILs in the field in Cologne, Germany. The parental accessions (mean days till flowering: PI 558499, 71.4 days; PI 604581, 114 days) and the assessed RILs showed high variability in flowering time (min. 53 days, max. 160 days; Figure 5A). The QTL mapping based on over 230,000 SNPs revealed two significant QTL regions, one on chromosome 10 (11,911,867 bp to 12,485,518 bp, 44 genes; one LOD range) and one on chromosome 6 (20,322,569 bp to 24,290,483 bp, 382 genes; one LOD range; Figure 5B, S7, Table S7). Both QTL showed additive genotypic effects (Figure 5C, S8), with the genotype of the early flowering parent PI 558499 leading to 16.4 and 11.5 days earlier flowering date than the population mean at the chromosome 10 and chromosome 6 QTL, respectively. Together, the two QTL result in a difference in flowering time of over 55 days between homozygous genotypes and account for a large proportion of the variation observed in this cross.

**Figure 5.**
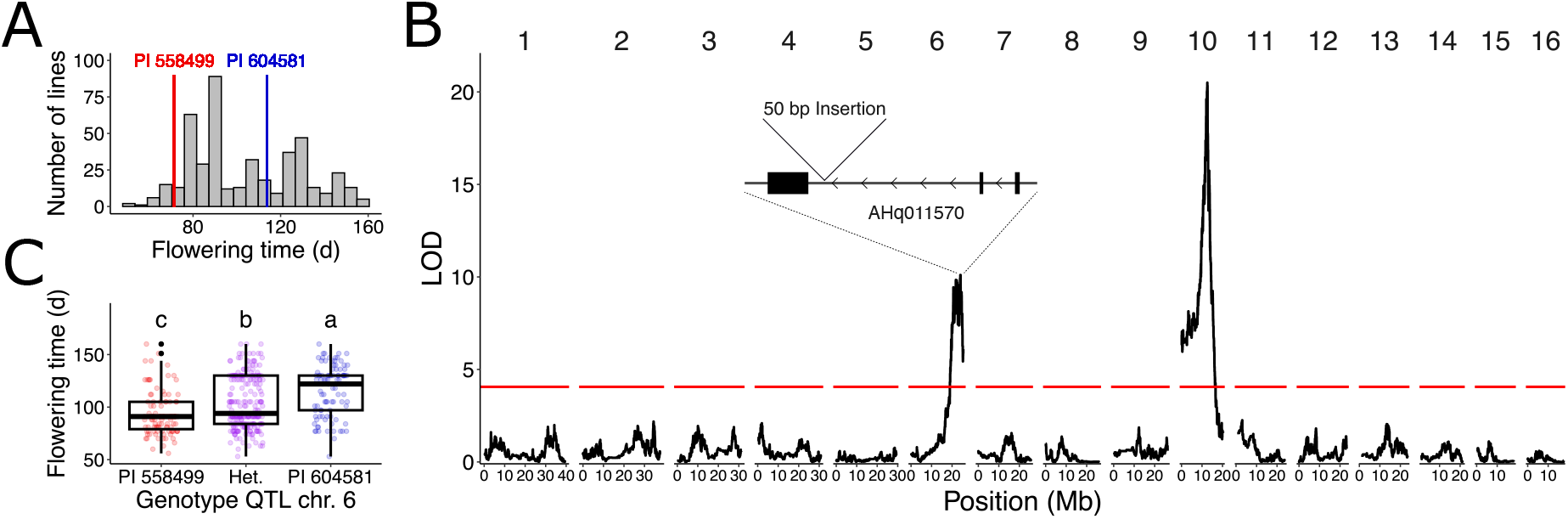
QTL mapping of flowering time in the *A. hypochondriacus* mapping population. (A) Flowering time distribution of F_3_ lines. The flowering times of the parental accessions (early flowering PI 558499 in red and late flowering PI 604581 in blue) as means from five replicates are indicated as vertical lines. (B) QTL mapping results for flowering time. The red line indicates the 95% LOD significance threshold based on 100 permutations. The gene model of the candidate gene AHq011570 within the chromosome 6 QTL is shown, including the position of the 50-bp insertion in the late-flowering parent PI 604581. Exons are depicted as black boxes, and arrows indicate the direction of transcription. (C) Flowering time for F_3_ lines based on the genotype at the QTL peak on chromosome 6. Differences in flowering time between genotypes were assessed using ANOVA and Tukey test and significant differences (p<0.05) between groups were displayed using compact letter display. Points are colored by genotype as in A, with heterozygotes in purple.

While the QTL regions included several genes, we identified candidate genes in the QTL regions based on their functional annotation. The QTL on chromosome 10 included an ortholog (AHq016812) of the *Arabidopsis thaliana* AP2 family transcription factor *TOE2* ∼35 kb from the QTL peak. *TOE2* functions as a floral repressor and represents a target of *miRNA 172* in the control of developmental timing (Aukerman and Sakai 2003). Due to the conserved function of *TOE2* in flowering time regulation and its position in the QTL region, AHq016812 represents a strong candidate gene for the control of flowering time in amaranth.

Within the QTL on chromosome 6, we identified multiple candidate genes, including AHq011814, an ortholog of *A. thaliana* flowering locus T located 560 kb from the QTL peak, and AHq011570, which encodes a CCCH-type zinc finger protein (KHZ-like) located approximately 2 Mb from the QTL peak. AHq011570 is orthologous to *KHZ1* in *A. thaliana*, a known regulator of flowering time via modulation of *FLC* expression (Yan et al. 2017). We found AHq011814 to be identical in protein sequence between the two parental accessions and AHq011570 to carry a single amino acid substitution (AHq011570 position 100, D to Y). However, an additional assembly and annotation of the late flowering parent (PI 604581) using Oxford Nanopore MinIon revealed structural variants associated with the identified candidate genes. The early flowering parent PI 558499 had a transposable element insertion in the promoter region approximately 2 kb (2,042 bp) upstream of AHq011814 and the late flowering PI 604581 contained a 50 bp insertion in the second intron of AHq011570 compared to the early flowering parent. Comparing gene expression across organs in PI 558499 revealed an over 300 times higher expression of AHq011814 in the flower than in the leaf Figure S9. The candidate genes and their associated structural variants are very promising candidates controlling natural variation in flowering time and make excellent targets for breeding efforts.

## Discussion

The grain amaranths represent a case of repeated, independent domestication from a shared wild ancestor, creating a species complex. Here, we present a pangenome for the grain amaranth species complex, composed of chromosome-level assemblies for the five species of the grain amaranth species complex. The pangenome provides a robust foundation for comparative and evolutionary analyses and enables a detailed examination of the genomic structural diversity associated with domestication. Our assemblies highlight the efficiency of modern plant genome assembly approaches. While the previous individual reference integrated multiple sequencing technologies — including PacBio HiFi, Oxford Nanopore, Hi-C, and Iso-Seq — our assemblies reach similar levels of contiguity and completeness using PacBio HiFi data alone (Tables 1 & S2; Graf et al. 2025). Compared to earlier assemblies of *A. cruentus* (Ma et al. 2021) and *A. hybridus* (Montgomery et al. 2020), our genomes show clear improvements in completeness and accuracy. Although gap-free reference genomes have been achieved for model crops, (Chen et al. 2023), these assemblies rely on extensive and costly multi-platform data integration, which remains a considerable barrier for minor crops. Nevertheless, our assemblies exceed the per-base accuracy of the T2T maize reference sequence, with QVs above 59 compared to a QV of 42.3 for maize (Chen *et al*. 2023, Table 1). These results demonstrate that even our individual assemblies for the grain amaranth species complex are comparable to high-quality genomes of major crops. While assemblies already existed for *A. hybridus, A. hypochondriacus* and *A. cruentus*, we provide the first assemblies for the South American grain species *A. caudatus* and its close relative *A. quitensis*. The highly complete genomes for all five species will serve as resource for breeding and functional studies within individual species.

The consistent assembly of all five species in the complex enables the direct comparison of their genome architecture and functional spaces. Our pangenome analysis revealed a conserved overall chromosome structure and gene content, suggesting that domestication of the grain amaranths was not driven by large-scale chromosomal restructuring or extensive gene gain or loss. Other crops experienced more extensive rearrangements in the course of domestication and selection, fixing important trait combinations (Jayakodi et al. 2024; Guo et al. 2025). Overall, the chromosome architecture showed a one-to-one correspondence between homologous chromosomes across the grain amaranth complex with ∼75% of groups of orthologous genes conserved across all five species. This proportion of the core genes is high compared to other plants, where often only ∼35-45% of genes are categorized as core: 35.9% in soybean (*Glycine max*) and wild relatives (Liu et al. 2020), 41.6% in rice (*Oryza sativa*) and wild relatives (Guo et al. 2025), and 43.7% in *Brassica oleracea* (Guo et al. 2024). The high proportion of core genes shows the close relationship between members of the grain amaranth species complex, but the within-species diversity in individual species might still be high and reveal more variable genes. Including more individuals will decrease the proportion of core genes as more individuals are compared (Guo et al. 2025; Liu *et al*. 2020; Zhao et al. 2025). For example, studies in foxtail millet (He et al. 2023a) and rice (Guo et al. 2025) analyzed more than 100 genomes, and a study in peanut combined eight complete assemblies with resequencing data from 269 additional accessions (Zhao *et al*. 2025). These larger-scale comparisons yield lower core gene percentages (23.8% in millet; 41.6% in rice) than the six-way comparison presented in this study.

Within the core genes of amaranth, we find enrichment for specific functions where gene copies were lost during domestication, potentially due to high genomic redundancy. This phenomenon may arise from gene duplication and subsequent relaxed selective constraints leading to the eventual loss of redundant copies (Wolf and Koonin 2013; Clark 2023). Conversely, gene gains in the core set during domestication include protein biosynthesis genes (Figure 4), suggesting that duplications may have contributed to the high protein content of the crop (Joshi et al. 2018).

A group of genes that is often associated with the variable genome are NLR genes. Across plants, NLR gene counts vary widely, from a single gene in *Utricularia gibba* to more than 1,000 in *Triticum aestivum* (Barragan and Weigel 2021). Grain amaranths have a moderate number of NLR genes (Figure S6), comparable to *Brassica rapa* (187) and *Fragaria vesca* (165) (Barragan and Weigel 2021). While the total number in each genome was similar between amaranth species, individual copies and clusters differed (Table S5, Table S6). The difference in number of NLR genes might result from different pathogen distribution in North and South America and individual NLR copies might provide resistance to local pathogens. These genes should be further studied through molecular assays to identify their specificity.

The comparison of gene sets offers insight into functional changes during domestication, and assessing a larger number of accessions per species will be essential to fully characterizing the diversity of the grain amaranth species complex. The high-quality reference genomes presented here will serve as a framework for these broader analyses.

Despite the high collinearity between genomes and the similar number of genes, amaranth has a high diversity of structural variation of different sizes (Figure 3). In total, we identified over 100,000 unique SVs by comparing our five new assemblies and the *A. hypochondriacus* v3 reference genome to the *A. retroflexus* reference genome (Raiyemo et al. 2025). This was substantially more than the ∼27,000 SVs a study found comparing 115 cucumber (*Cucumis sativus*) accessions (Zhang *et al*. 2015), or the ∼25,000 between 347 Sorghum (*Sorghum bicolor*) genotypes (Songsomboon et al. 2021). In grain amaranth, SVs are often unique to a single species and are not consistently shared across all three domesticated species (Figure 2). This pattern further supports the repeated, independent domestication of the three grain amaranths that has also been found with SNP-based population genetic analyses (Stetter et al. 2020; Gonçalves-Dias et al. 2023). Hence, SVs reflect the domestication history of the crops. These unique rearrangements contribute to genomic diversity and could be linked to species divergence and the domestication process.

Domesticate-specific SVs may underlie phenotypic differences relevant to domestication, as SVs are often the causal variants of domestication-related traits (Gaut et al. 2018). This has been suggested for one-third of known domestication genes (Gaut et al. 2018). Here, we map the genetic control of an essential life history trait, flowering time and find two major QTL controlling the trait. Both include important flowering time regulators, known from model plants. Two promising candidate genes in the QTL on chromosome 6 were associated with SVs in potential regulatory regions in the promoter and introns (Figure 5). One of of these candidate genes (AHq011570) differs between the parents by a 50-bp insertion, making it a potential candidate underlying the substantial difference in flowering time observed in the population (Figure 5). The transposon insertion in the promoter region of flowering locus T (AHq011814) may alter the regulation of the gene and influence flowering. Structural variation associated with flowering locus T has been shown to alter flowering time in sunflower and has been associated with habitat adaptation (Todesco et al. 2020). Such structural variation could easily be missed by genotyping using short-read sequencing data, demonstrating the value of the pangenome for the discovery of SVs potentially causal for phenotypic differences. Further molecular work in an isogenic background is needed to validate the candidate genes and to assess the impact of SVs on these genes and their expression patterns.

## Conclusions

Here, we present a genomic resource for the entire grain amaranth species complex, particularly in the context of domestication. We assembled five high-quality, chromosome-scale genomes for each of the five species in the complex, including improved assemblies for *A. hybridus* and *A. cruentus* and the first assemblies for *A. caudatus* and *A. quitensis*. The merged pangenome provides a strong foundation for genomics studies in amaranth and enabled us to observe whole-genome architecture changes across the entire species complex and to examine the potential role of SVs in amaranth speciation and domestication. Differences in gene content between wild and domesticated species showed similar functional enrichment patterns, suggesting that domestication reshaped existing pathways — such as photosynthesis, protein metabolism and RNA regulation — rather than creating entirely new functions. We identify two QTL driving a 55-day difference in flowering time, likely partially controlled by structural variation. Overall, this work establishes both a high-quality genomic framework and new insights into grain amaranth domestication, providing a platform for future studies that will expand our understanding of the genomic basis of domestication and adaptation.

## Methods

### Plant material and sequencing

We identified a representative accession from each of the five species in the grain amaranth species complex for genome sequencing and assembly based on short read sequencing information and geographic origin (Figure 1). These data were obtained from the amaranth genetics and genomics database (AmaranthGDB: https://amaranthgdb.org/; Gonçalves-Dias and Stetter 2021). We extracted high molecular weight (HMW) DNA from leaves of a single plant of PI 511754 (*A. hybridus*), PI 643036 (*A. hypochondriacus*), PI 643058 (*A. cruentus*), PI 642741 (*A. caudatus*), and PI 490466 (*A. quitensis*), using the Macherey-Nagel Nucleobind HMW DNA extraction kit (Macherey-Nagel GmbH & Co KG, Düren, Germany) following manufacturer’s instructions. The DNA was sequenced using PacBio HiFi long-read sequencing with Novogene (Novogene GmbH, Planegg, Germany).

### Genome assembly and annotation

We converted the sequencing data from binary alignment map (BAM) format to FastQ format using the “bam2fastq” option from the pbtk Bioconda toolkit v3.1.1 (PacBio 2023), and ran NanoPlot v1.42.0 and NanoQC v0.10.0 from the NanoPack2 suite of tools (De Coster and Rademakers 2023) to ensure that no adapter sequences were present and that sequencing reads were of high accuracy (quality value (QV) >20). We then assembled nuclear genomes from PacBio HiFi reads using Hifiasm v0.25.0 (Cheng et al. 2021, 2022, 2024) with default parameters. Primary contigs were converted from GFA to FASTA format using *awk*.

Chloroplast and mitochondrial genomes were assembled from the same HiFi reads using OATK v1.0 (Zhou et al. 2025), with the coverage threshold (-c) set to 100 to exclude most nuclear reads, as whole-genome sequencing coverage was below 100x. We annotated the chloroplast genomes using the web-based GeSeq v2.03 (Tillich et al. 2017) with default settings, using the NCBI RefSeq from *A. hypochondriacus* and the MPI-MP Reference Set “chloroplast land plants (CDS + rRNA)” as references for BLAT searches, and ARAGORN v1.2.38 (Laslett and Canback 2004) for tRNA annotation. Mitochondrial genomes were annotated using the PMGA webtool (Li et al. 2025) with the “29 Mitogenomes” reference database. Gene fragments with <60% coverage were inspected and manually removed from the annotations if the fragment covered <60% of the reference gene. Remaining partial matches were classified as pseudogenes based on evaluation with ORF Finder (https://www.ncbi.nlm.nih.gov/orffinder/) and BLAST (https://blast.ncbi.nlm.nih.gov/Blast.cgi), as they did not encode full-length protein products and were therefore not counted as genes. Chloroplast and mitochondrial genome structures and annotations were visualized using OG-Draw (Greiner et al. 2019).

To remove organellar genome sequences in the nuclear assemblies, we mapped the nuclear contigs to the assembled chloroplast and mitochondrial genomes using Minimap2 v2.28 (Li 2018, 2021). Scaffolds and contigs showing >90% sequence identity over >70% of their length were removed from the nuclear assembly. The filtered nuclear contigs were then scaffolded against the *A. hypochondriacus* v3 reference genome of PI 558499 (Graf et al. 2025) with the “scaffold” function in RagTag v2.1.0 (Alonge et al. 2022), which does not break contigs, thereby preserving true structural differences between assemblies. Structural errors were identified and corrected using Inspector v1.3 (Chen et al. 2021) in reference-free mode. We combined the final nuclear assembly with the organellar assemblies into a single assembly FASTA per species.

After assembly and organellar sequence filtering, we annotated transposable Elements (TEs) and other repetitive sequences with EDTA v2.2.2 (Ou et al. 2019). These annotations were used to soft-mask repetitive regions to reduce false gene predictions. Structural gene annotation was performed with Helixer v0.3.4 (Holst et al. 2023) on the soft-masked assemblies. The resulting GFF output was processed with gffread v0.12.8 (Pertea and Pertea 2020) to generate GTF files and to extract coding sequences (CDS) and protein sequences in FASTA format. Gene models lacking a valid start and/or stop codon were removed from the gene annotation. We evaluated annotation quality using BUSCO v5.8.2 (Manni et al. 2021b,a; Li 2023) in “proteome” mode using the “embryophyta_odb10” lineage file, and generated summary statistics using AGAT v1.5.0 (Dainat et al. 2024). Functional annotation of protein sequences was performed using the web-based tools EggNOG-mapper v2.0.1 (Cantalapiedra et al. 2021; Huerta-Cepas et al. 2019) and Mercator4 v7.0 (Bolger et al. 2021; Schwacke et al. 2019).

For each of the assembled genomes we detected telomeric repeats, using the “search” function in tidk v0.2.65 (Brown *et al*. 2025), with the The query string (-s) set to “AAACCCT,” the canonical telomere repeat in *Amaranthus* (Lightfoot et al. 2017; Peska and Garcia 2020), and a window size (-w) of 1,000 bp. Contig boundaries were similarly identified using tidk “search,” this time with the query string set to “N,” corresponding to the 100 Ns inserted between contigs during RagTag scaffolding. Telomere positions and contig breakpoints were used to guide limited manual curation of assemblies. In cases where Rag-Tag scaffolding had clearly inverted terminal regions and local contiguity supported confident correction, terminal segments spanning from chromosome ends to telomeric repeat arrays were inverted to restore telomere placement (before assembly annotation). We used CentIER v2.0 (Xu *et al*. 2024) with default parameters to annotate centromere positions, using the gene annotation GFF file as an additional input.

Genome quality and completeness were evaluated using BBMap v39.10 (Bushnell 2022), QUAST v5.3.0 (Gurevich *et al*. 2013), LAI (Ou et al. 2018), and Merqury v1.3 with Meryl v1.4.1 (Rhie et al. 2020).

### Synteny and structural variation

We performed whole-genome interchromosomal synteny analyses with NGenomeSyn v1.43 (He *et al*. 2023b) to identify and visualize syntenic blocks between genome assemblies. NGenomeSyn was run using Minimap2 as the mapper, with minimum length set to 250kb. We also analyzed intrachromosomal synteny with SyRI v1.7.0 (Goel et al. 2019). We aligned the assemblies using Minimap2 v2.28 (Li 2018, 2021), identified syntenic and non-syntenic regions — including SVs — with SyRI (Goel et al. 2019), and visualized the results with plotsr (Goel and Schneeberger 2022).

To identify and compare SVs across genomes, all five assemblies and the *A. hypochondriacus* v3 reference genome were first aligned to the *A. retroflexus* reference genome (Raiyemo *et al*. 2025) using Minimap2 v2.28 (Li 2018, 2021). Next, we used SVIM-asm v1.0.3 (Heller and Vingron 2021) to call SVs from the Minimap2 alignments with minimum size set to 50 bp, and merged these into one Variant Call Format (VCF) file with the “merge” function from SURVIVOR v1.0.7 (Jeffares et al. 2017) with minimum supporting caller adjusted to one to keep all unique SVs. SVs were merged irrespective of strand, but only if they were of the same type and within 1,000 bp of each other. Comparisons between species and plotting was done in R v4.3.1 (R Core Team 2023).

### Orthologous gene identification

We used OrthoFinder v3.1.0 (Emms and Kelly 2019; Emms *et al*. 2025) to identify orthologous genes between the five new assemblies and the *A. hypochondriacus* v3 reference genome. To improve accuracy, we included *Beta vulgaris* (Dohm et al. 2014) as an outgroup. For downstream gene content comparisons and orthogroup-based statistics, we used hierarchical orthogroups inferred at node N1, which refers to the internal node of the species tree representing all six amaranth genomes but excluding *B. vulgaris*. We analyzed the overlap in between genomes to identify core genes (those in orthogroups with at least one gene from all assemblies), and those unique to certain groups or individual species and plotted them with the UpsetR package (Conway et al. 2017; Lex et al. 2014). We also identified orthogroups that, among the core orthogroups, gained or lost genes in the domesticated species (*A. hypochondriacus, A. cruentus*, and *A. caudatus*) relative to their wild ancestor (*A. hybridus*). We subsequently did an enrichment analysis with Mercator4 v7.0 (Bolger et al. 2021; Schwacke et al. 2019) on these genes to determine whether they are significantly enriched in any particular functional categories. We used NLRtracker v1.0.3 (Kourelis et al. 2021) to identify nucleotide-binding leucine-rich repeat (NLR) genes in the grain amaranth pangenome, using the protein FASTA file for each genome assembly as input.

### Linkage mapping of flowering time

To investigate the genetic control of flowering time, we used a segregating population generated from the *A. hypochondriacus* accessions PI 558499 (Plainsman) and PI 604581 (Winkler *et al*. 2026). For the linkage mapping, we used imputed genotype calls of the F_2_ generation for 230,124 bi-allelic SNPs differentially fixed between the two parental accessions (Winkler *et al*. 2026). We grew F_3_ offspring lines and five replicates of both parental accessions in a randomized field trial in Cologne, Germany, between April and October 2025, and recorded flowering time. We performed F_2:3_ QTL mapping using R/qtl2 (Broman et al. 2019), defining QTL based on 95% significance thresholds from 100 permutations. We implemented an iterative mapping approach using a custom script, adding the genotypes with maximum marginal probability at the highest QTL peak as covariates to identify additional QTL (Arthur et al. 2025). To estimate genotype effects on flowering time, we calculated Best Linear Unbiased Predictors (BLUP) including the kinship matrix to account for residual polygenic effect as implemented in R/qtl2 (Broman et al. 2019).

The late flowering parent (PI 604581) was sequenced on an Oxford Nanopore MinIon. Reads were filtered with chopper v0.9.2 from Nanopack2 (De Coster and Rademakers 2023) to retain only those with quality >Q10, yielding ∼18x coverage and a mean read length of 6,236 bp. Filtered reads were assembled with Flye v2.9.3 (Kolmogorov et al. 2019) and scaffolded against the published *A. hypochondriacus* v3 reference genome of PI 558499 (Graf et al. 2025) without breaking contigs. The resulting assembly was 401.2 Mb in length, composed of 161 scaffolds, with a scaffold N50 of 24.8 Mb. Annotation was performed with EDTA and Helixer as described for the five PacBio HiFi assemblies (see Genome assembly and annotation). We called SVs between the two parents as described previously, using early-flowering parent (PI 558499, *A. hypochondriacus* v3) as the reference. Gene-SV overlaps were identified using a custom script. The gene expression of candidate genes was evaluated in expression data values from Graf *et al*. (2025).

### Data visualization

Unless otherwise stated, all statistical analysis and data visualization was done in RStudio (Posit team 2023) using R v4.3.1 (R Core Team 2023). Packages used include: circlize (Gu *et al*. 2014), data.table (Barrett et al. 2024), GenomicRanges (Lawrence et al. 2013), ggbreak (Shuangbin Xu et al. 2021), ggpattern (Fc et al. 2024), ggplot2 (Wickham 2016), maps (Becker et al. 2024), RColorBrewer (Neuwirth 2022), tidyverse (Wickham et al. 2019), and UpSetR (Conway et al. 2017; Lex et al. 2014).

## Declarations

### Ethics approval and consent to participate

Not applicable.

### Consent for publication

Not applicable.

### Availability of data and materials

All raw sequencing data have been deposited in the European Nucleotide Archive (ENA) at EMBL-EBI under accession number PRJEB108719 for the long-read data used for the genome assemblies and under accession number PRJEB106936 for the short-read data used for QTL mapping. Genome assembly and annotation files can also be accessed on ENA under accession number PRJEB108719 and on AmaranthGDB https://amaranthgdb.org/. Custom scripts used for all analyses in this study are available on GitHub at https://github.com/cropevolution/PanAma1_2026.

### Authors’ contributions

M.G.S. conceptualized, designed and supervised the project. E.L. developed the genome assembly pipeline, performed comparative genomics analyses, and drafted the manuscript. T.S.W. phenotyped plants and performed the QTL mapping and data analysis. E.L., T.S.W., and M.G.S. interpreted the results and edited the manuscript. All authors read and approved the final manuscript.

### Authors’ information

Institute for Plant Sciences and Cluster of Excellence on Plant Sciences, University of Cologne, Cologne, 50674, Germany

Ella Ludwig, Tom S Winkler, Markus G Stetter

## Acknowledgments

We would like to thank Evelin Fahle and Roswitha Lentz for plant care and DNA extractions. We also thank Corbinian Graf for developing the QTL mapping population and Sophie Feyerabend for help with phenotyping in the field trial.

## Supplementary Information

**Figure S1.**
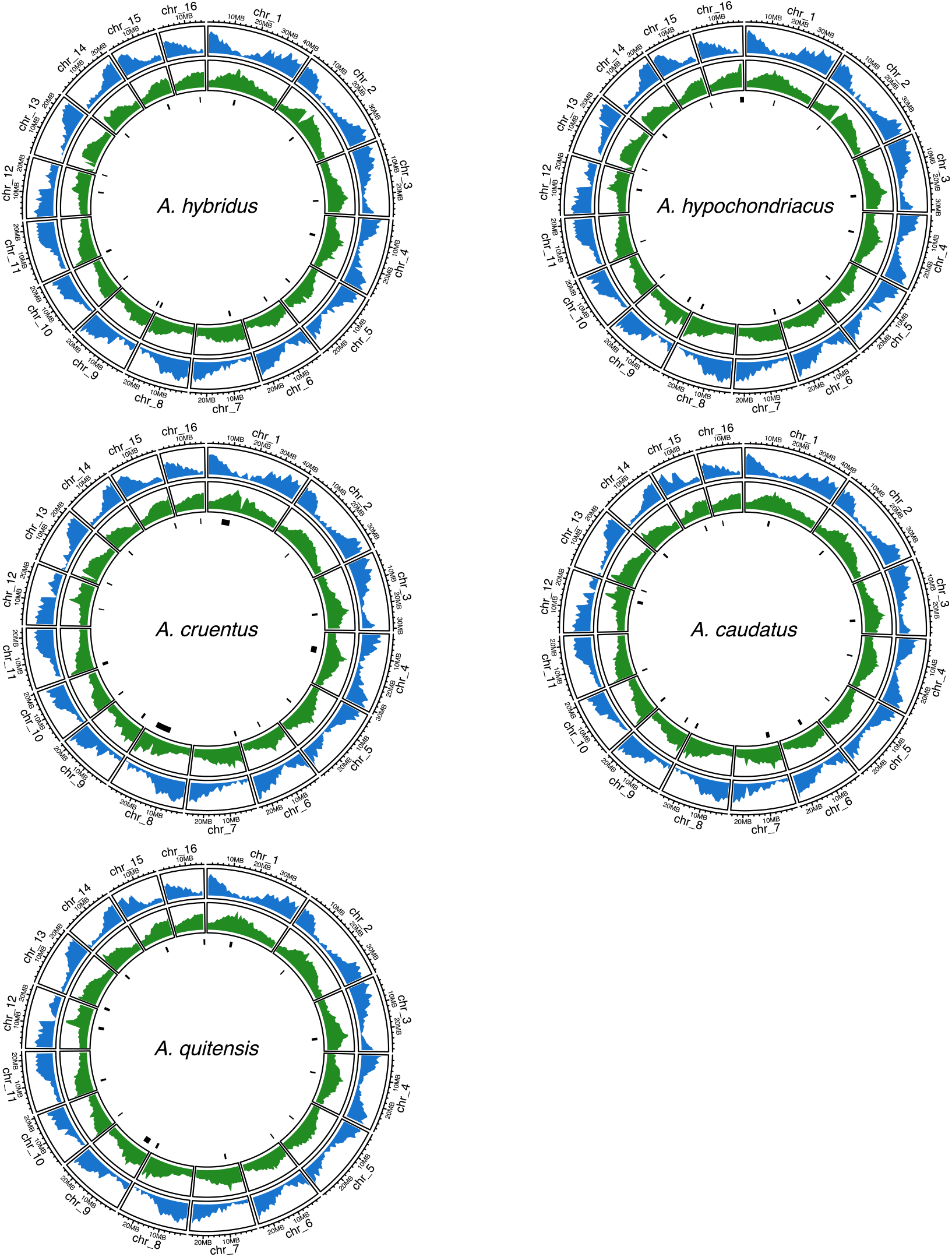
Genomic distribution of genes and TEs in the five new amaranth genome assemblies. Tracks depict the distribution of annotated genes (blue) and TEs (green), calculated in 1 Mb windows, and predicted centromeres (black bars).

**Figure S2.**
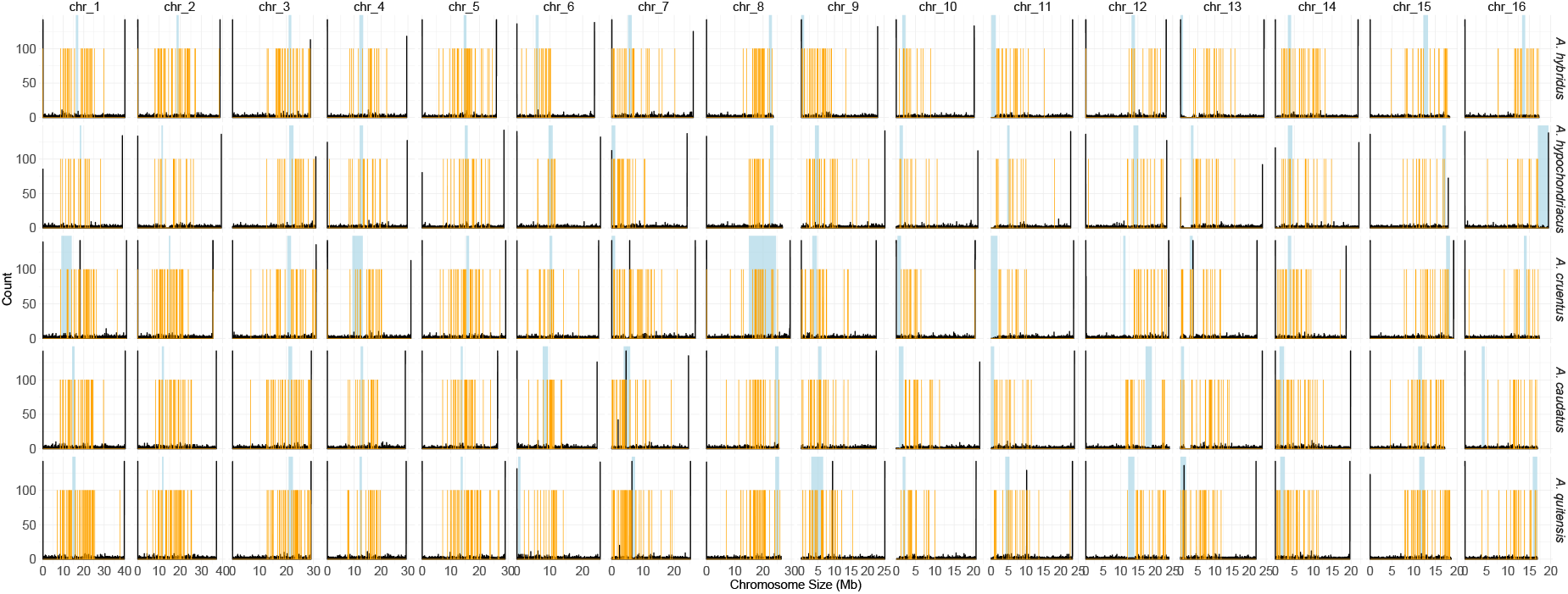
By-chromosome distribution of telomeric repeats, predicted centromeres, and contig breaks across the five amaranth genomes. Telomeric repeats, counted in 1 kb windows, are shown in black, predicted centromere ranges are indicated by blue shading, and contig breaks are marked in orange.

**Figure S3.**
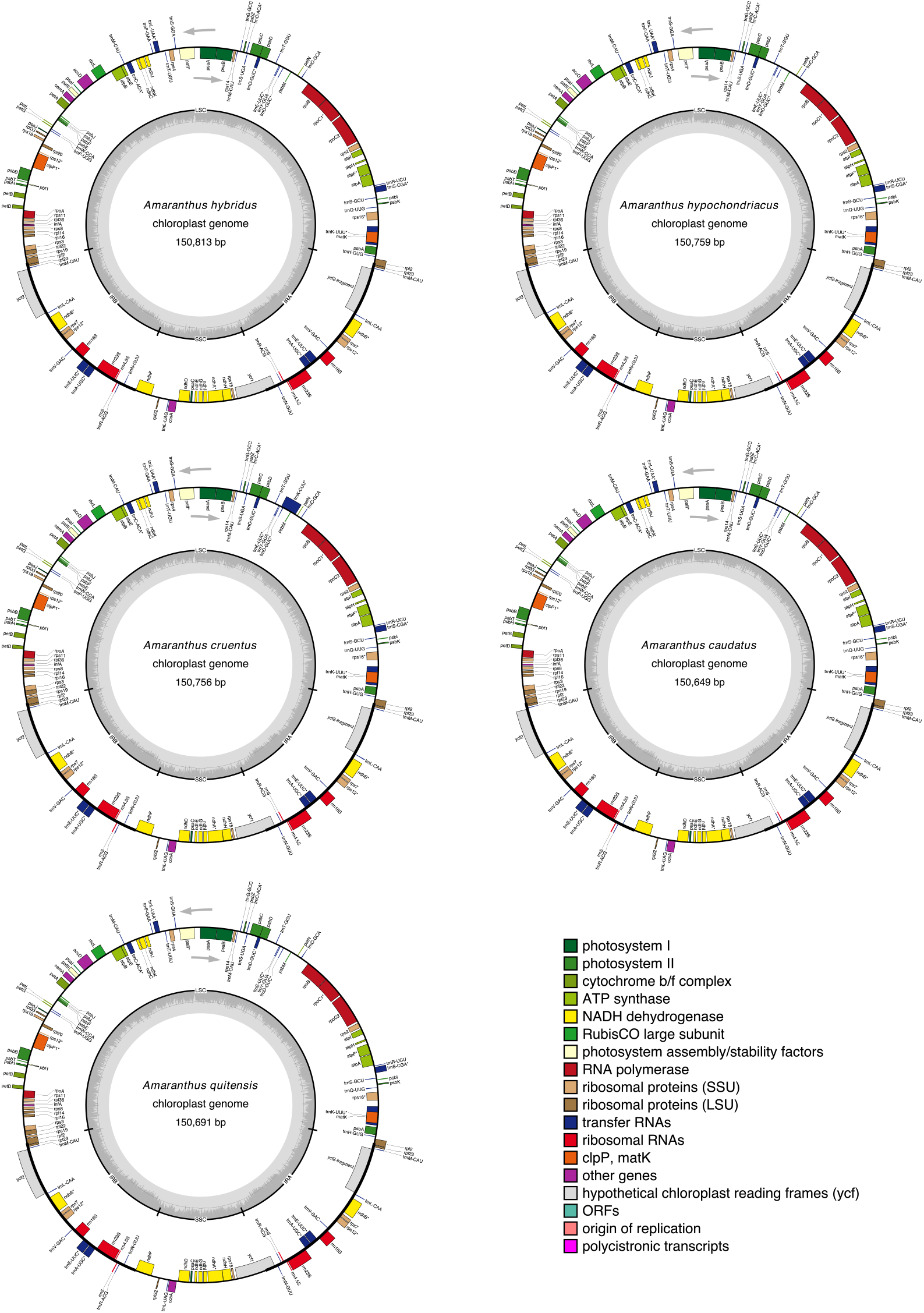
Structures and gene annotations of chloroplast genomes of five amaranth species. Annotated genes are shown in the outer tracks, colored according to functional category, with genes containing introns marked with *. Arrows indicate the direction of transcription. GC content is plotted in the central track. The quadripartite structure of the chloroplast genome is shown, with the large single-copy (LSC) region, small single-copy (SSC) region, and two inverted repeats (IRA, IRB) marked.

**Figure S4.**
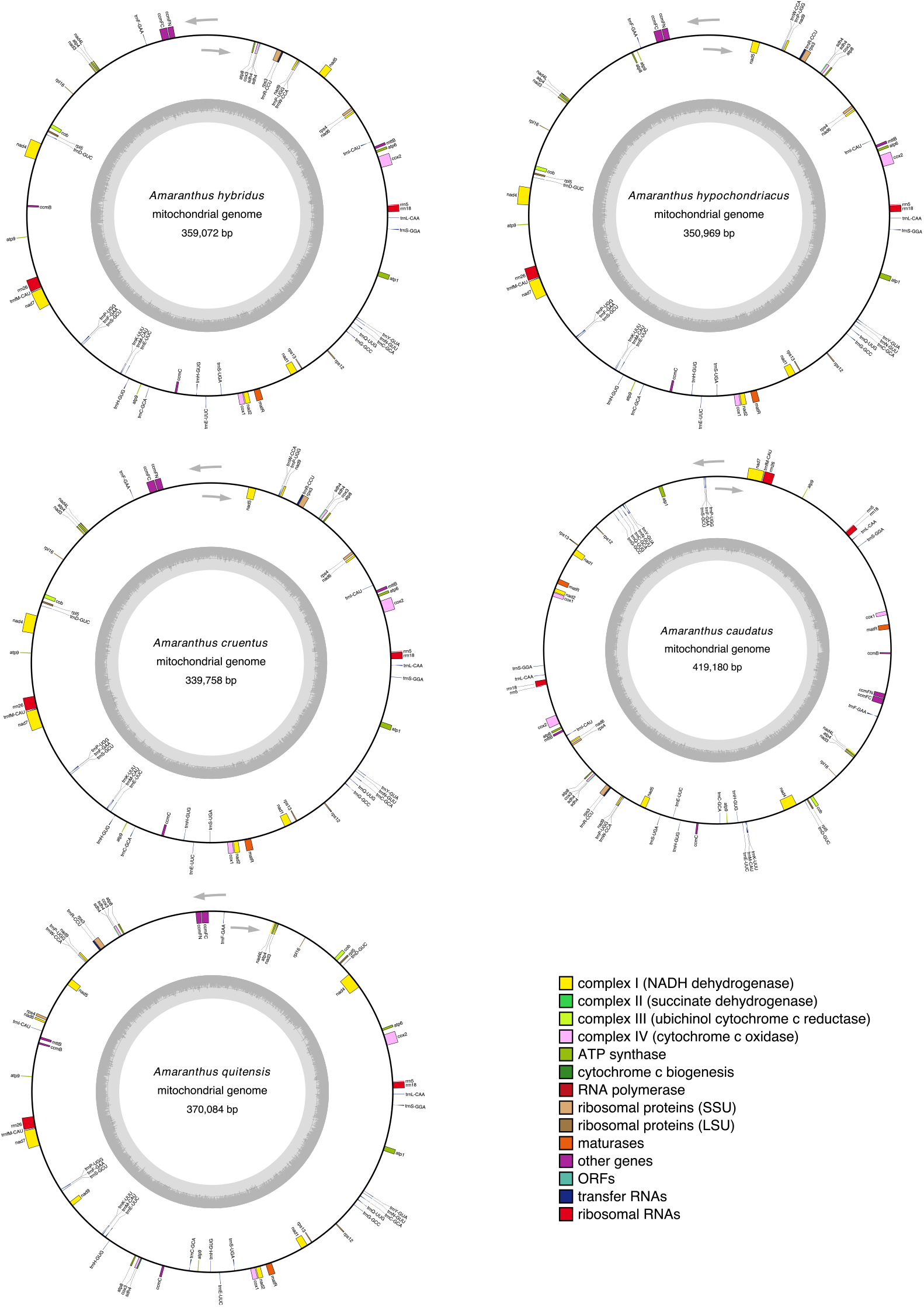
Structures and gene annotations of mitochondrial genomes of five amaranth species. Annotated genes are shown in the outer tracks, colored according to functional category. Arrows indicate the direction of transcription. GC content is plotted in the central track.

**Figure S5.**
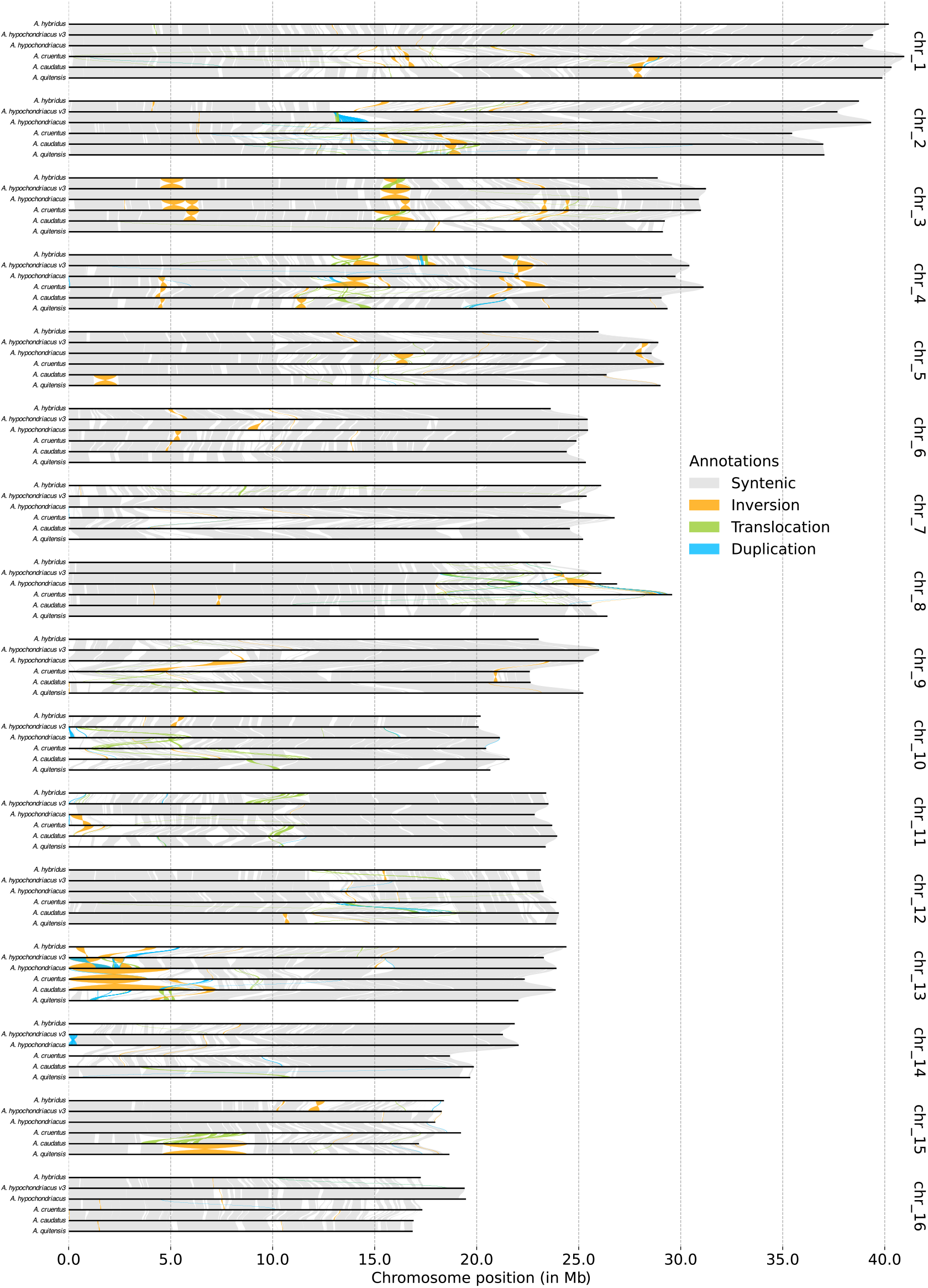
Intrachromosomal synteny of all chromosomes among grain amaranth species complex. Syntenic blocks are shown in gray, inversions in orange, translocations in green, and duplications in blue.

**Figure S6.**
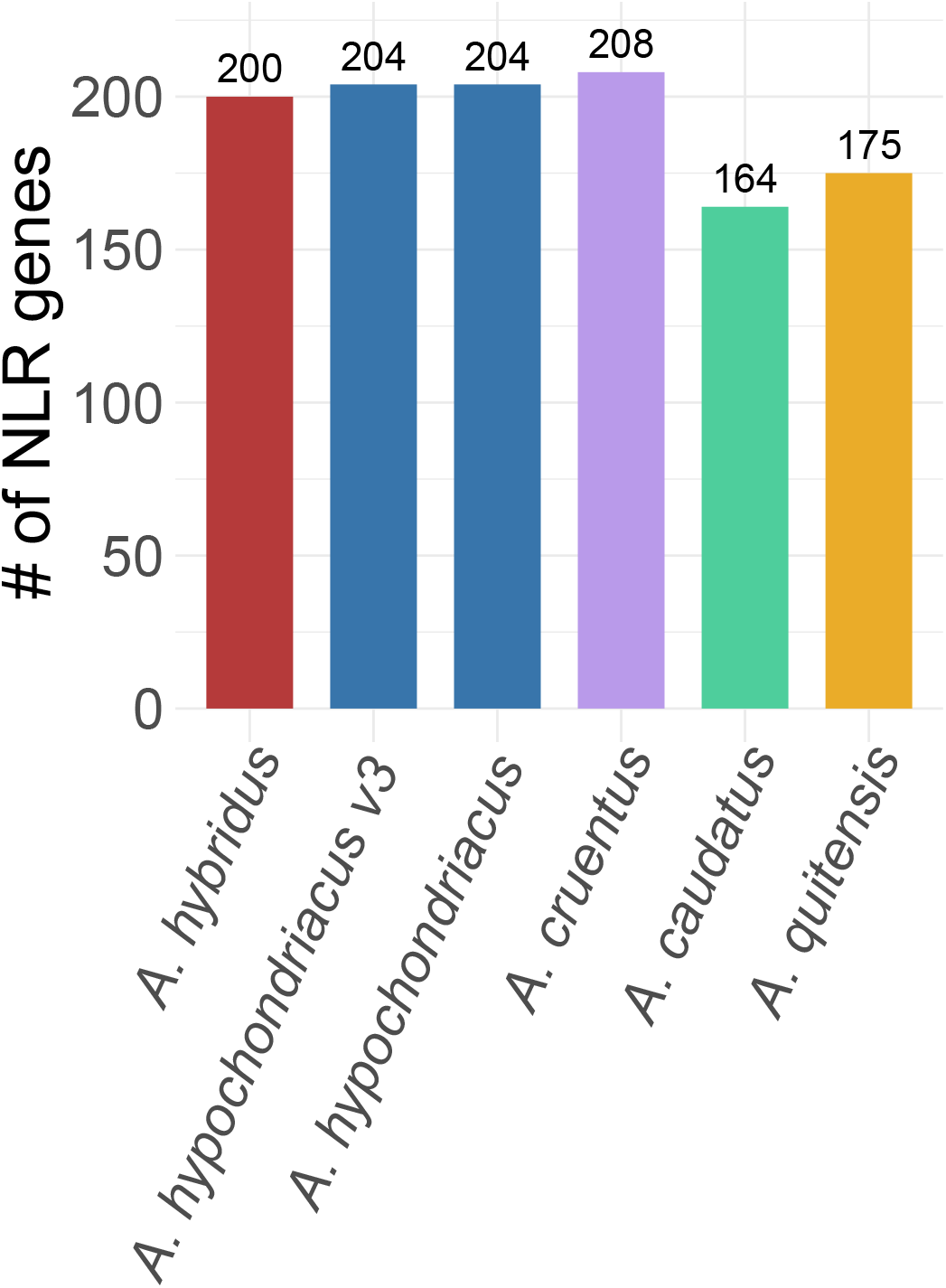
Number of NLR genes predicted per assembly. Bars are colored by species.

**Figure S7.**
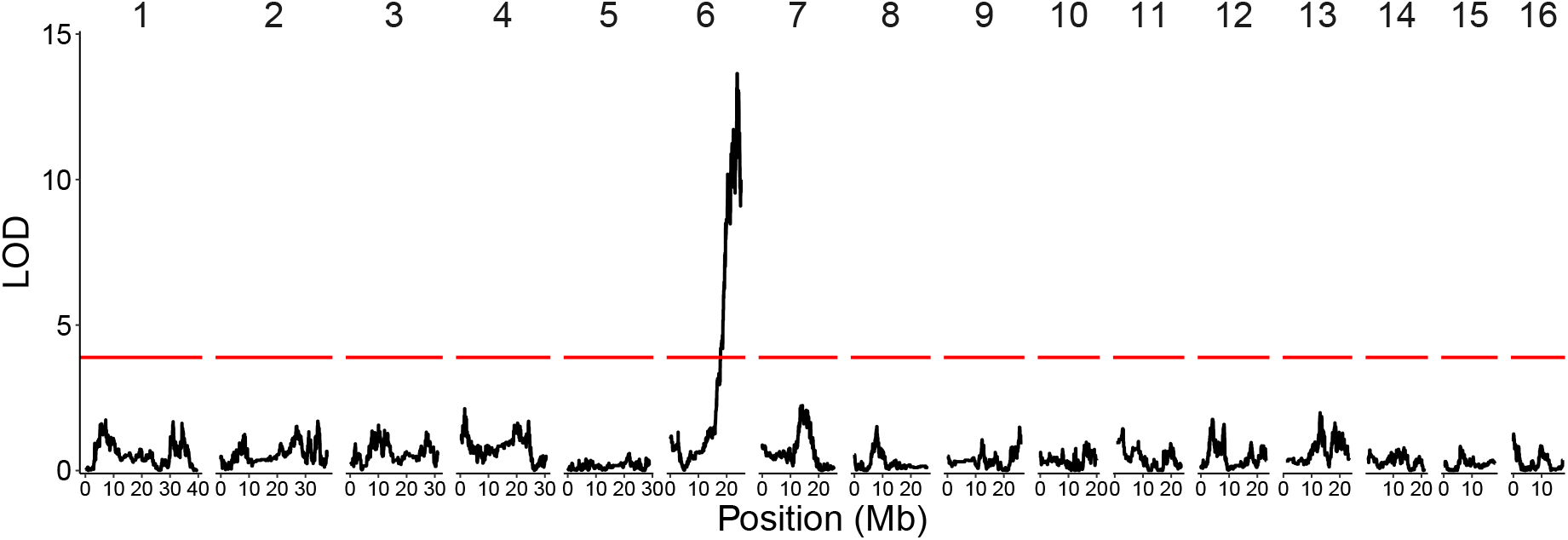
QTL mapping for flowering time using RIL genotypes at the first detected QTL on chromosome 10 as covariates. The red line depicts the 95% LOD significance threshold based on 100 permutations.

**Figure S8.**
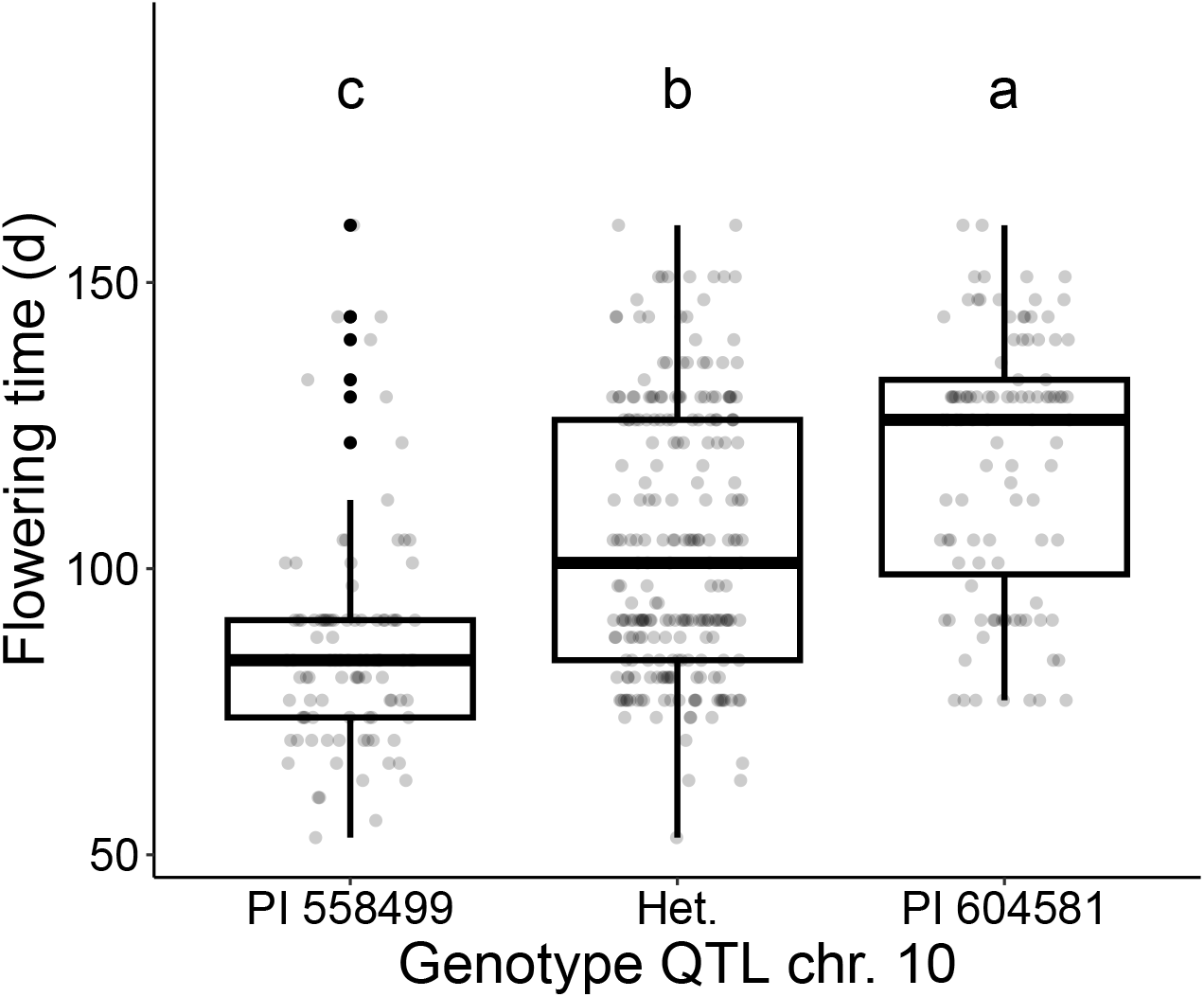
Association between flowering time and genotype at the chromosome 10 QTL peak for F_3_ lines. Differences in flowering time between genotypes were assessed using ANOVA and Tukey test and significant differences (p<0.05) between groups were displayed using compact letter display.

**Figure S9.**
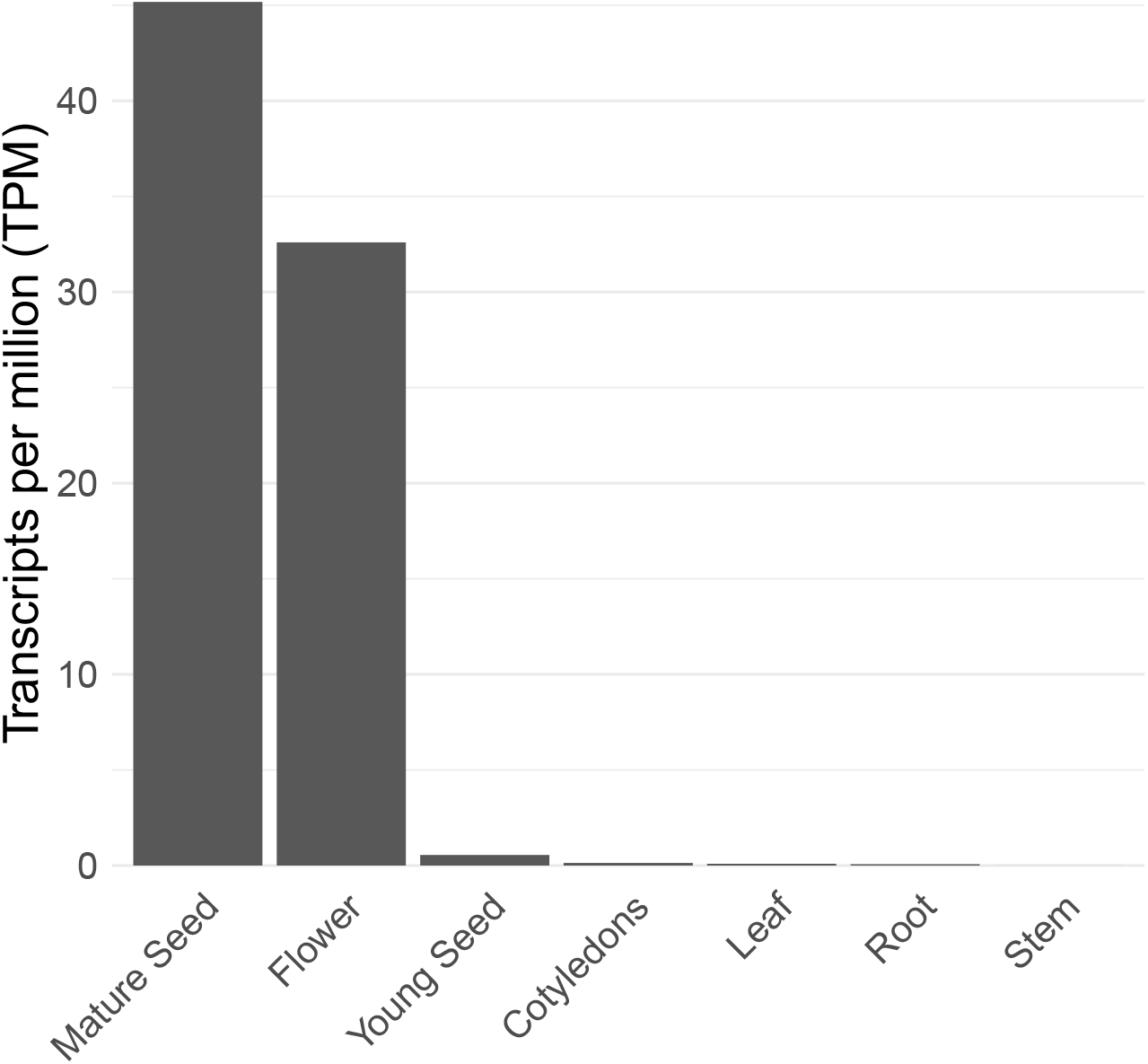
Expression (in transcripts per million) of AHq011814 in different tissues in PI 558499.

**Table S1.** PacBio HiFi long-read sequencing data statistics. Sequencing coverage was calculated by dividing the total number of bases by the estimated genome sizes from Stetter and Schmid (2017) provided in Table 1.

|  | *A. hybridus* | *A. hypochondriacus* | *A. cruentus* | *A. caudatus* | *A. quitensis* |
| --- | --- | --- | --- | --- | --- |
| Mean read length (bp) | 17,931 | 16,539 | 16,760 | 11,842 | 12,308 |
| Mean read quality | 26.6 | 24.4 | 26.8 | 30.0 | 28.7 |
| Number of reads | 1,906,687 | 1,016,607 | 2,014,448 | 2,088,062 | 1,512,241 |
| Total bases (Gb) | 34.2 | 16.8 | 33.8 | 24.7 | 18.6 |
| Sequencing coverage | 68x | 33x | 67x | 49x | 37x |

**Table S2.**
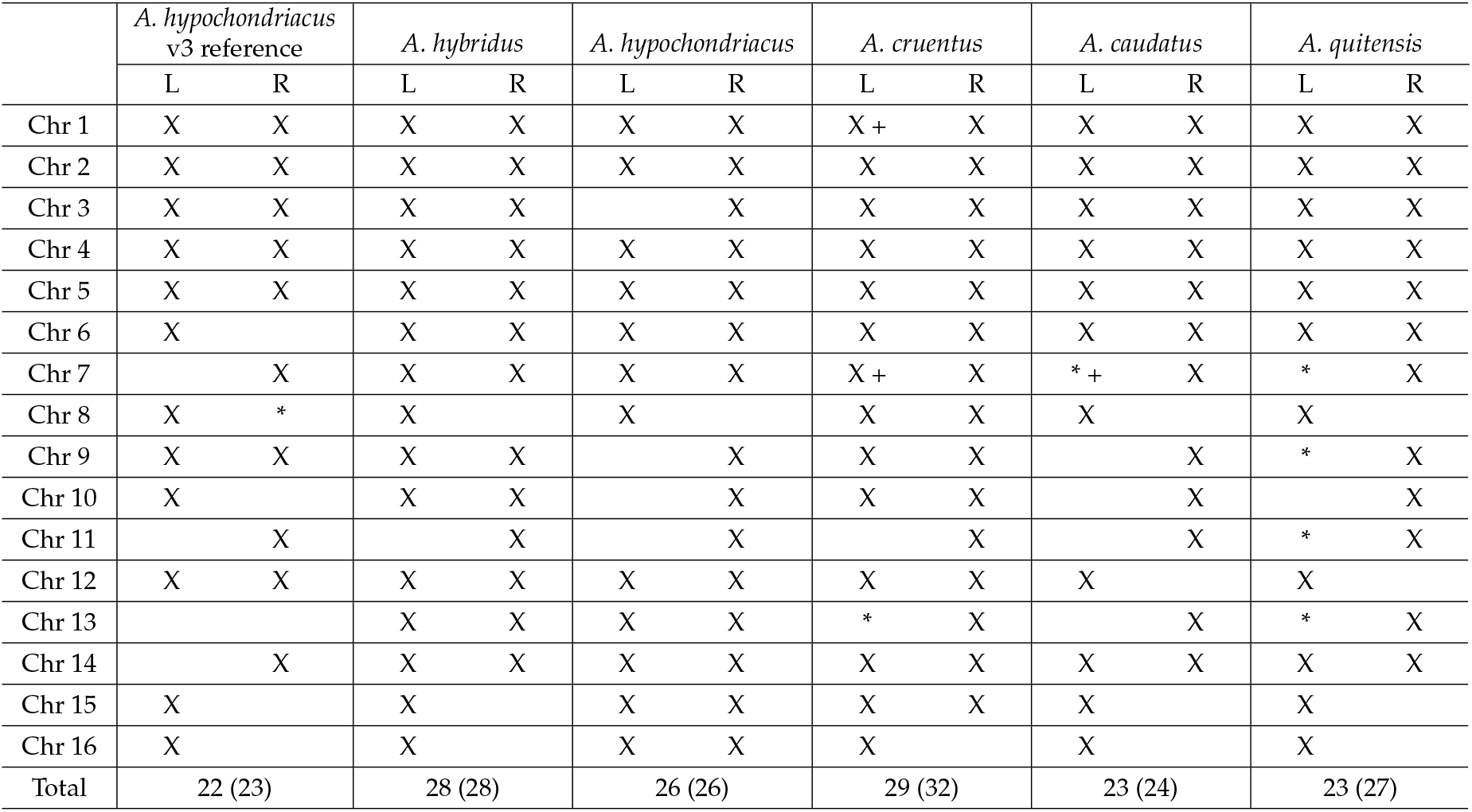
Telomeres present in the five new grain amaranth genome assemblies and the published *A. hypochondriacus* v3 reference. An ‘X’ indicates that the telomere was identified and correctly assembled at the end of the chromosome, while ‘*’ indicates a telomere not at the end of the chromosome and ‘+’ denotes the presence of a third telomere on a chromosome. In the “Total” row, the first number is the number of telomeres at the ends of chromosomes (assumed to be correctly assembled), and the number in parentheses, denotes the total number of telomeres detected, whether at the end of the chromosomes or not.

**Table S3.** Assembly and annotation statistics for chloroplast and mitochondrial genomes of five grain amaranth species.

|  | *A. hybridus* | *A. hypochondriacus* | *A. cruentus* | *A. caudatus* | *A. quitensis* |
| --- | --- | --- | --- | --- | --- |
| **Chloroplast genome** |  |  |  |  |  |
| Assembly length (bp) | 150,813 | 150,759 | 150,756 | 150,649 | 150,691 |
| Length of inverted repeat (IR) | 23,538 | 23,538 | 23,538 | 23,528 | 23,528 |
| Length of large single copy (LSC) | 85,770 | 85,729 | 85,726 | 85,636 | 85,687 |
| Length of small single copy (SSC) | 17,967 | 17,954 | 17,954 | 17,957 | 17,948 |
| Number of genes | 126 | 126 | 127 | 126 | 126 |
| Number of protein-coding genes | 79 | 79 | 79 | 79 | 79 |
| Number of tRNAs | 39 | 39 | 40 | 39 | 39 |
| Number of rRNAs | 8 | 8 | 8 | 8 | 8 |
| GC content | 36.6% | 36.6% | 36.6% | 36.6% | 36.6% |
| **Mitochondrial genome** |  |  |  |  |  |
| Assembly length (bp) | 359,072 | 350,969 | 339,758 | 419,180 | 370,084 |
| Number of genes | 62 | 61 | 61 | 62 | 62 |
| Number of protein-coding genes | 34 | 33 | 33 | 34 | 34 |
| Number of tRNAs | 25 | 25 | 25 | 25 | 25 |
| Number of rRNAs | 3 | 3 | 3 | 3 | 3 |
| GC content | 42.7% | 42.8% | 42.8% | 42.8% | 42.6% |

**Table S4** Orthogroups of annotated genes identified by OrthoFinder (Emms and Kelly 2019) across all assemblies. Each row corresponds to an orthogroup, and columns list the member genes from each assembly, enabling identification of orthologous genes across genomes. Orthologs were inferred using *B. vulgaris* as an outgroup. Shown here are both the full set of orthogroups (including *B. vulgaris*) and hierarchical orthogroups inferred at node N1, representing the internal node of the species tree that includes all six amaranth genomes but excludes *B. vulgaris*.

**Table S5** Predicted nucleotide-binding leucine-rich repeat (NLR genes) and their functional annotations for five new grain amaranth genome assemblies and the previously published *A. hypochondriacus* v3 reference genome (Graf et al. 2025).

**Table S6** Orthogroups of nucleotide-binding leucine-rich repeat (NLR) genes identified by OrthoFinder across all assemblies. Each row corresponds to an orthogroup, and columns list the member genes from each assembly, enabling identification of orthologous NLR genes across genomes.

**Table S7** List of genes in *A. hypochondriacus* flowering time QTL. Genes in the two flowering time QTL are listed along their chromosomal position and functional annotation based on Graf *et al*. (2025). The functional annotation for all isoforms include the general description, preferred name, PFAMs, *A. thaliana* orthologs, mercator annotations, and all assigned gene ontology terms for each isoform.

